# Analog closed-loop optogenetic modulation of hippocampal pyramidal cells dissociates gamma frequency and amplitude

**DOI:** 10.1101/279588

**Authors:** Elizabeth Nicholson, Dmitry A Kuzmin, Marco Leite, Thomas E Akam, Dimitri M Kullmann

## Abstract

Gamma-band oscillations are implicated in modulation of attention and integration of sensory information. The finding that cross-regional coherence varies with task and performance suggests a role for gamma oscillations in flexible communication among anatomically connected brain areas. How networks become entrained is incompletely understood. Specifically, it is unclear how the spectral and temporal characteristics of network oscillations can be altered on rapid timescales needed for efficient communication. We use closed-loop optogenetic modulation of principal cell excitability to interrogate the dynamical properties of hippocampal oscillations. Gamma frequency and amplitude can be modulated bi-directionally, and dissociated, by phase-advancing or delaying optogenetic feedback to pyramidal cells. Closed-loop modulation alters the synchrony rather than average frequency of action potentials, in principle avoiding disruption of population rate-coding of information. Modulation of phasic excitatory currents in principal neurons is sufficient to manipulate oscillations, suggesting that feed-forward excitation of pyramidal cells has an important role in determining oscillatory dynamics and the ability of networks to couple with one another.

## Introduction

Gamma-band (approximately 30 to 120 Hz) oscillations have been implicated in the modulation of attention and perception, in action initiation, spatial navigation and memory encoding, and have also been proposed to underlie flexible information routing among anatomically connected regions^1–9^. Central to several of these proposed roles is the ability of gamma oscillations in different areas to enter into, and exit, states of synchrony with one another^10–12^. Evidence for behavioral-state dependent coupling and uncoupling comes from variable oscillatory coherence among distinct components of the visual cortex, correlating with selective stimulus attention^13,14^. An earlier study in the rodent hippocampal formation showed that the CA1 subfield can flip between a state of coherence with the medial entorhinal cortex at ~110 Hz and a state of coherence with the CA3 subfield at ~40 Hz, correlating with information flow through the temporo-ammonic and Schaffer collateral pathways respectively^15^. Although several experimental confounds cloud the interpretation of coherence measured from local field potential (LFP) recordings^16^, these studies provide some of the most compelling evidence that gamma-band oscillatory entrainment underlies flexible functional connectivity.

Although the cellular mechanisms underlying gamma oscillations have been extensively studied^17,18^, there remain uncertainties over the fundamental determinants of their dynamics and the relative contributions of excitatory and inhibitory signaling. Gamma-band oscillations can be induced *in vitro* in the presence of blockers of ionotropic glutamate receptors^19^, or *in vivo* by optogenetic stimulation of parvalbumin-positive interneurons^20,21^, underlining the importance of fast perisomatic inhibition^22–24^. Robust population oscillations can also be simulated in exclusively inhibitory networks^25^. These experimental and computational observations emphasize the importance of inhibitory kinetics. Nevertheless, gamma-band oscillations can be entrained by sinusoidal optogenetic stimulation of pyramidal neurons in an *in vitro* hippocampal slice preparation^10^. This observation implies that phasic depolarization of principal cells can determine the gamma rhythm and argues against a model where the only role of pyramidal cells is to tonically depolarize a network of reciprocally coupled interneurons^17,26^.

Further insight into the dynamical mechanisms of synchronization between oscillating networks comes from examining the phase response curve (PRC) of the network oscillation, defined as the phase advance or delay produced by a transient stimulation, as a function of the instantaneous phase at which the stimulus is delivered. The finding that gamma in an *in vitro* hippocampal slice preparation shows a biphasic PRC^10^ is consistent with the hypothesis that this oscillation can be entrained by appropriately modulated afferent activity. The shape of the PRC is furthermore accurately reproduced with a simple neural mass model^27^, where extracellular electrical or optogenetic stimuli are represented as transient perturbations of the instantaneous level of excitation or inhibition^10^. Recent theoretical work has derived population phase response curves for oscillations in spiking network models, providing insight into how mechanisms of oscillation generation determine entrainment properties^28,29^. Nevertheless, there remains a large gap between the PRC and understanding the determinants of the oscillatory frequency and interactions between gamma-generating circuits.

The present study investigates the dynamical properties of gamma oscillations by using closed-loop optogenetics to create an artificial feedback loop between the oscillatory network activity (as assessed by the LFP) and excitatory input to the principal cell population. Specifically, we delivered analog-modulated excitation whose strength was a function of the instantaneous phase and amplitude of the oscillation. This approach is quite distinct from previous closed-loop applications of optogenetics^30^, which have adopted one of four main strategies. First, several studies have used the detection of a change in the state of a network, such as the onset of an electrographic seizure^31,32^ or sharp-wave ripple^33^, to trigger light delivery and return the network to its ground state. Second, light pulses have been timed according to the phase of a theta oscillation^34^, while examining the consequences for behavior. In the latter example the theta oscillation itself was not altered. Third, optogenetics has been used to regulate the overall activity of a population of neurons at a desired level ^35^. Fourth, optogenetic depolarization of interneurons, triggered by spikes in an individual principal cell, has been used to simulate a feedback inhibitory loop to interrogate their role in gamma^21,36^. The goal of the present investigation is qualitatively different: to understand how the spectral characteristics of gamma are affected by rhythmic excitation arriving at different phases. Computational simulations have suggested that closed loop optogenetics could be used to adjust the phase of gamma^37^, but whether it can alter its frequency or amplitude remains unclear.

## Results

### Closed-loop feedback modulation of affects gamma oscillations in CA1

We expressed the red-shifted optogenetic actuator C1V1^38^ in the mouse hippocampus CA1 under the *Camk2a* promoter to bias expression to excitatory neurons. The local field potential (LFP) was recorded in the CA1 pyramidal cell layer in acute hippocampal slices. A slowly increasing ramp of light (peak wavelength 590 nm) was delivered via a light-emitting diode (LED) coupled to the epifluorescence port of an upright microscope, eliciting a gamma oscillation (Fig. 1a, b), as previously reported in rodents^10,39–42^, cats^43^ and monkeys^44^.

**Fig. 1.**
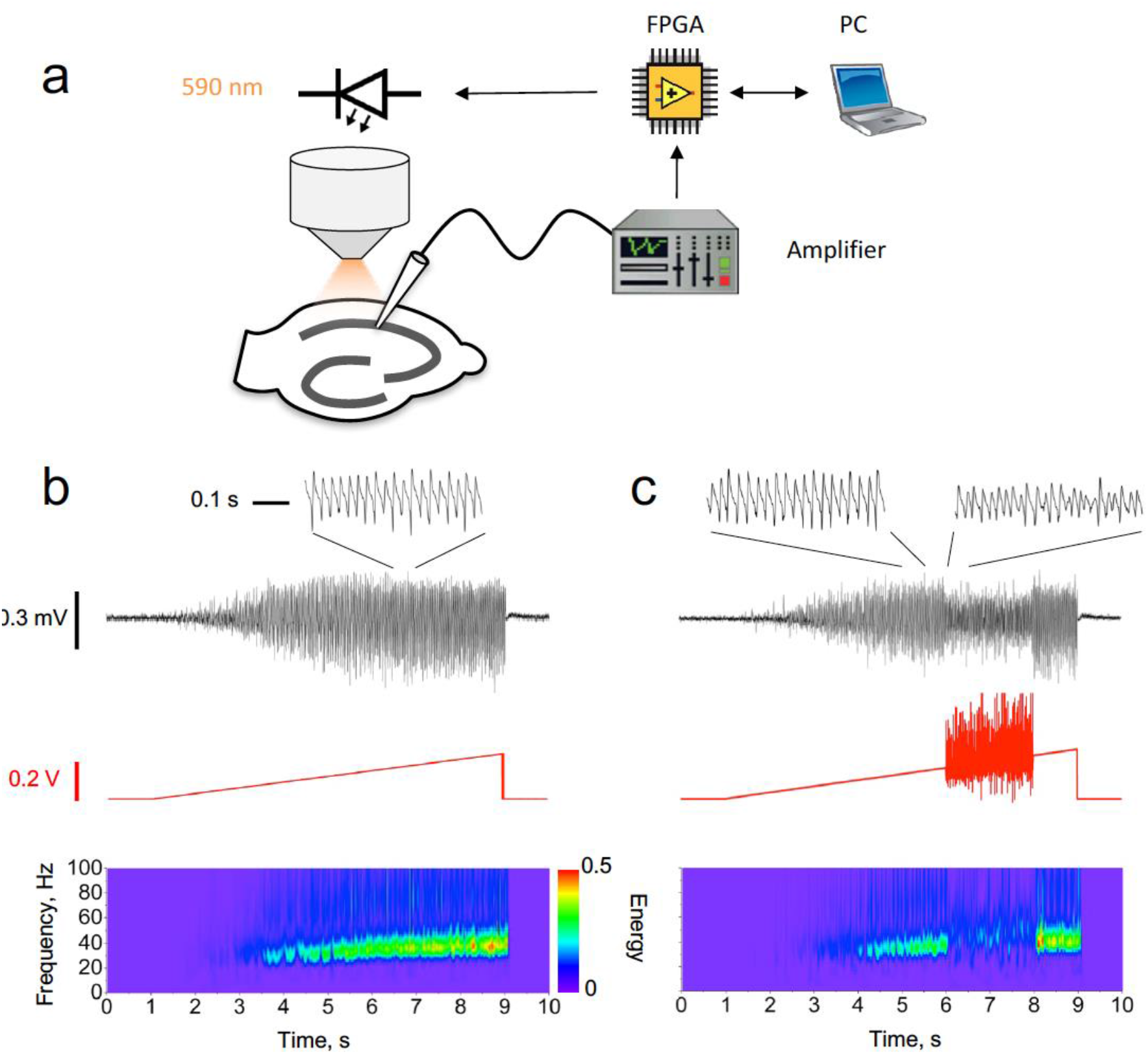
Closed-loop modulation of gamma oscillation. **(a)** Experimental design. The LFP in CA1 was used to modulate a ramp command generated by the PC via a field-programmable gate array (FPGA). The modulated ramp voltage command was then passed to the light-emitting diode (LED) driver, which implemented a threshold-linear voltage-to-current conversion. **(b)** Unmodulated oscillation recorded in CA1 induced by a linear ramp LED driver command. Black trace: LFP with an expanded section showing the characteristic shape of the gamma oscillation (inset). Red trace: LED ramp command. Bottom: LFP Morlet wavelet spectrogram. **(c)** Closed-loop oscillation clamp applied between 6 and 8 s, obtained by multiplying the ramp command by (1 + *k*_1_LFP + *k*_2_*d*LFP/*d*t), with *d*LFP/*d*t averaged over 2 ms intervals. For this example, *k*_1_ = 0 mV^−1^, *k*_2_ = 25 ms mV^−1^. The oscillation amplitude was reduced by approximately 60% (insets), with no net change in frequency.

In order to investigate the role of phasic excitation in setting the dynamical properties of gamma we used the LFP itself to manipulate the optogenetic drive in real time. The LED driver command was multiplied by a simple function of the instantaneous value of the LFP and its time-derivative: (1 + *k*_1_ LFP + *k*_2_ *d*LFP/*d*t)), where *k*_1_ and *k*_2_ are positive or negative constants. These operations were implemented with a field-programmable gate array (FPGA) and applied for a defined duration (typically 1 or 2 seconds) during the ramp. This yielded a change in the spectral properties of the oscillation, which lasted for the duration of the closed-loop feedback (Fig. 1c). Because both the LFP and its time-derivative fluctuated about 0, the “gamma clamp” had little effect on the average illumination intensity relative to an unmodulated ramp. Changes in the oscillation frequency or power could therefore not be attributed to a net increase or decrease in the average optogenetic drive to pyramidal neurons.

We adjusted the clamp function by altering the values of *k*_1_ and *k*_2_, and asked whether the frequency and/or power of the gamma oscillation can be modulated bidirectionally. Changes in spectral properties were related to the phase difference between the LFP and the LED drive during the clamp, as estimated from the cross-spectrum at maximal magnitude. Because the LFP is typically non-sinusoidal^45^, the phase difference was approximate. In-phase modulation, achieved by setting *k*_1_ positive and *k*_2_ = 0, led to an increase in oscillatory power and frequency (Fig. 2a). Modulating the ramp in anti-phase relative to the LFP, by setting *k*_1_ negative, led to a decrease in both frequency and power (Fig. 2b). Advancing the phase of the clamp by approximately 90°, achieved by setting *k*_1_ = 0 and *k*_2_ positive, increased the frequency of the oscillation whilst decreasing is power (Fig. 2c). Finally, a decrease in frequency and increase in power was achieved by delaying the trough of the clamp modulation relative to the LFP, by setting *k*_2_ negative (Fig. 2D). Detailed inspection of the ramp command waveform during the clamp shows that it was in some cases distorted relative to the LFP (e.g. Fig. 2b, c).

**Fig. 2.**
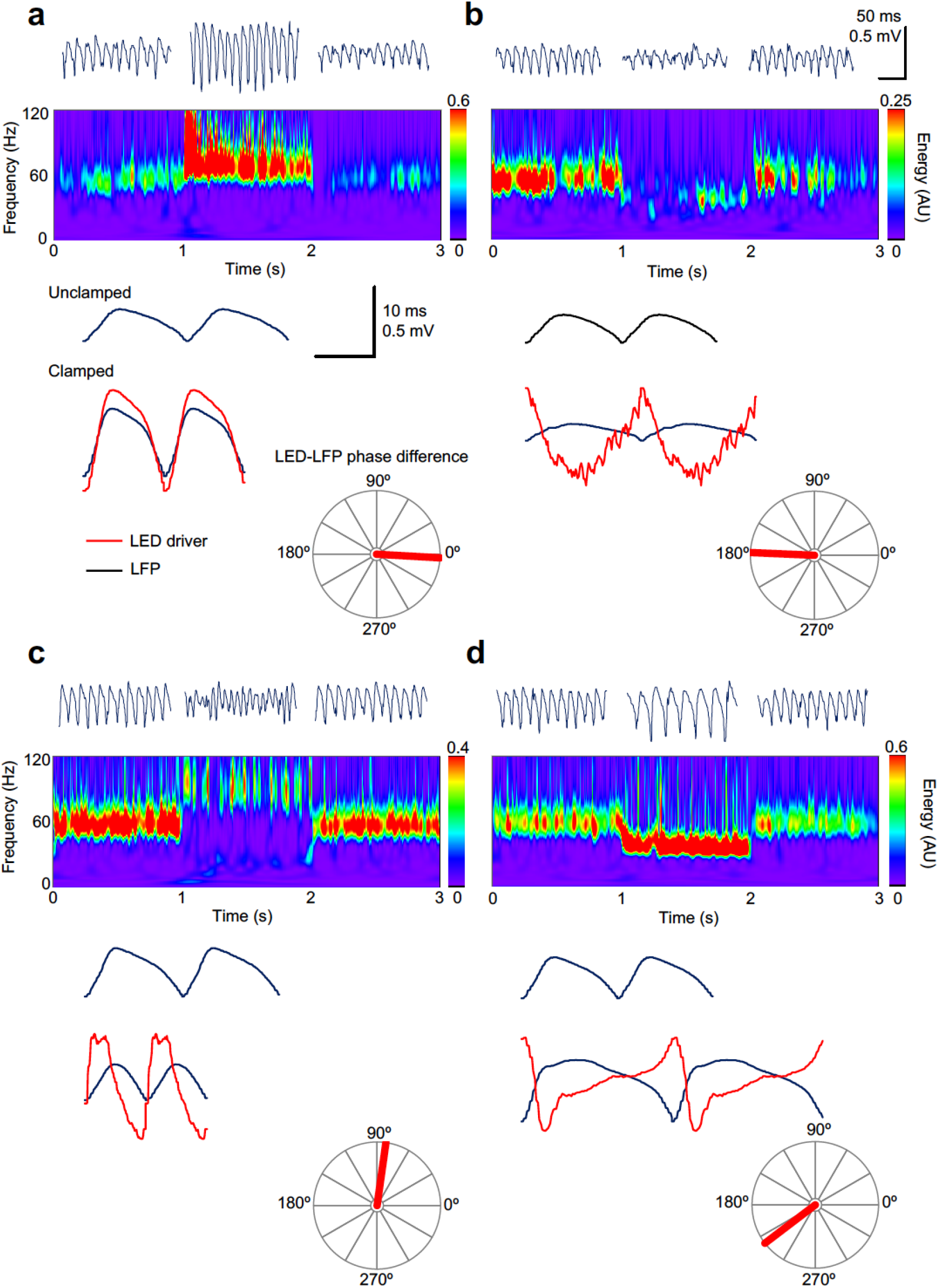
“Gamma clamp” allows bidirectional modulation of frequency and power. **(a)** In-phase modulation led to an increase in gamma power and frequency. Top: 200 ms-long segments of the LFP before, during and after closed-loop modulation of the LED driver. Middle: spectrogram. Bottom: two cycles of the average oscillation before and during the oscillation clamp. The average LED command (red trace, arbitrary scale) is shown superimposed on the clamped oscillation. The polar plot shows the phase relationship between the LED command and the LFP. **(b)** Anti-phase modulation led to decreases in both frequency and power. **(c)** An increase in oscillation frequency, together with a decrease in power, was obtained with ~90° phase-advance of the LED driver command over the LFP. **(d)** A decrease in frequency, together with an increase in power, was obtained when the LED modulation was delayed relative to the LFP by ~145 °. Scale bars apply to all panels.

Attempts to estimate the instantaneous oscillation phase, for instance using a Hilbert transform, and to use this to phase-advance or phase-delay a template of the LFP, compressed or stretched in time, were unsuccessful: the phase jitter and cycle-to-cycle variability in the amplitude and frequency of the gamma oscillation (see LFP traces in Fig. 2) prevented accurate estimation of these parameters in the face of closed loop feedback.

### Oscillation clamp is broadly consistent with the phase response curve of gamma

Changes in frequency and power, expressed in relation to the approximate phase difference between the LED command and the LFP, were qualitatively consistent across experiments (Fig. 3a-c). Moreover, as the LED-LFP phase difference was rotated through a complete cycle, the effect on the oscillation in the two-dimensional plane defined by the change in oscillation frequency and power also rotated through 360°, such that with the appropriate phase of closed-loop feedback the network oscillation could be pushed in any desired direction in the oscillation frequency-power space (Fig. 3c).

**Fig. 3.**
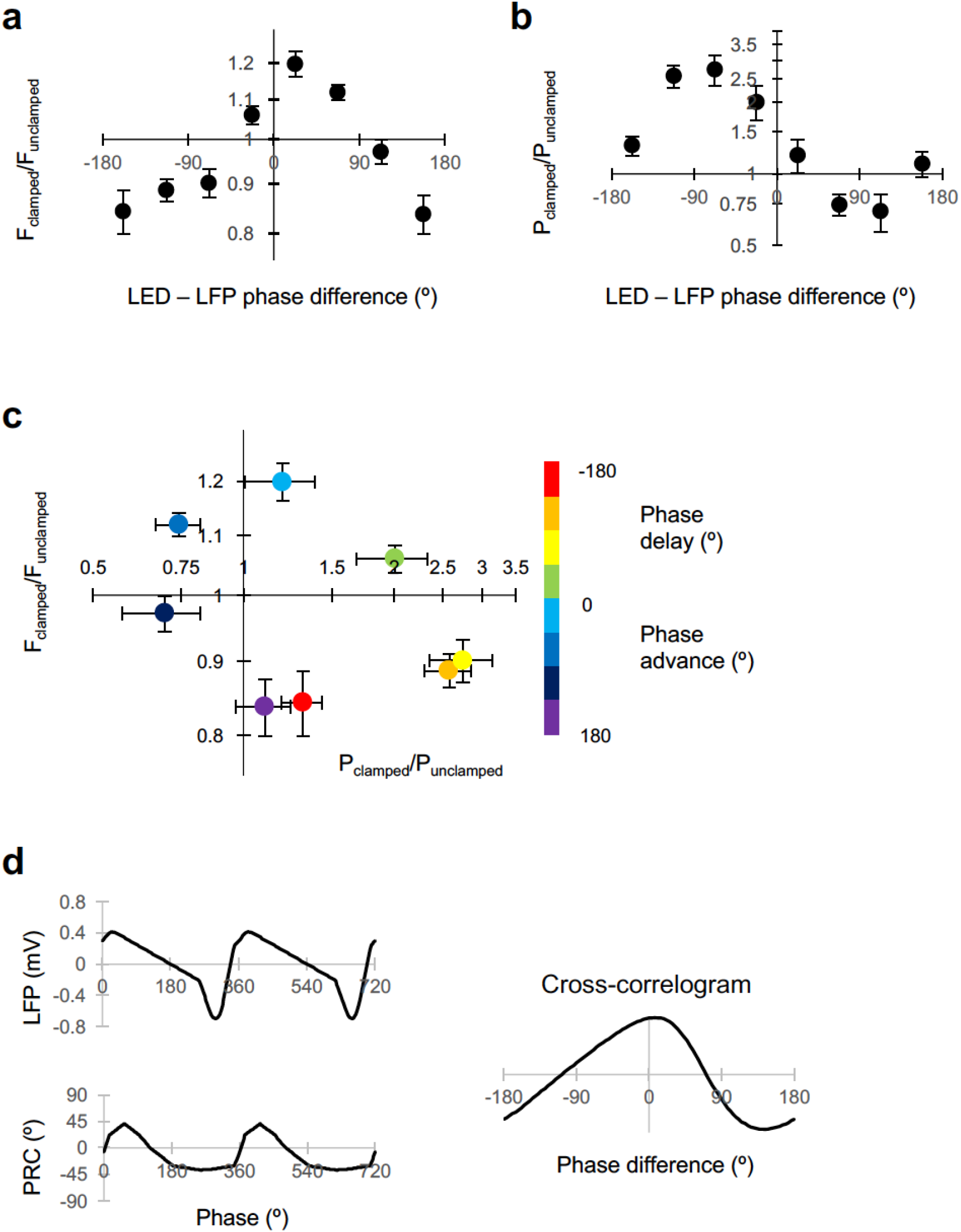
Dissociable modulation of oscillation frequency and power. **(a)** Dependence of frequency change on the phase relationship between the LED modulation and the LFP (positive values indicate LED phase advance relative to LFP). Changes in frequency are plotted as Fclamped/Funclamped, where the unclamped frequency was averaged from the gamma oscillation for 1 s before and 1 s after the gamma clamp was applied. Data are shown as mean ± SEM (n = 19 experiments). A positive phase difference indicates that the modulation was phase-advanced relative to the LFP. **(b)** Dependence of power change on the phase difference plotted as in (A). **(c)** Change in frequency plotted against change in power for different LED – LFP phase differences (color code at right). **(d)** Average LFP and phase response (PRC) curve from ^10^ (left). The circular cross-correlogram at right yields a prediction of the effect of a continuous modulation on the oscillation frequency, in rough agreement with the observed relationship in (A).

To gain a mechanistic insight, we asked if the characteristic relationship between the frequency change and the LED-LFP phase difference could be explained by the shape of the phase-response curve (PRC) previously reported^10^. In that study, a brief ‘kick’ was applied on top of the LED ramp command, and the phase advance or delay of subsequent oscillations was related to the phase of the LFP at which the transient occurred. A phase delay was observed when the transient optogenetic stimulus was delivered at the trough of the LFP, when pyramidal neurons are most likely to fire. The maximal phase advance, in contrast, occurred when the stimulus was delivered approximately one third of a cycle after the trough of the LFP. Assuming linear behavior, the effect of modulating the light intensity in closed loop can be obtained by averaging the product of the phase shift and the LFP over the entire cycle of the oscillation. The circular cross-correlogram between the typical LFP shape and the PRC should then predict the effect of modulating the optogenetic drive by the shape of the LFP itself at arbitrary degrees of phase advance or delay (Fig. 3d). In-phase modulation is expected, on the basis of this calculation, to phase-advance the oscillation, and thus to result in an increase in oscillatory frequency over several cycles. Anti-phase modulation, in contrast, is predicted to phase-delay the oscillation, and thus to decrease is frequency. The circular cross-correlation is, moreover, asymmetrical, broadly consistent with the shape of the relationship between the change in frequency and LED-LFP phase difference observed in the clamp experiments (Fig. 3a).

Although the shape of the PRC predicts the changes in gamma frequency achieved with closed loop modulation at different LED-LFP phase differences, on its own it says nothing about changes in power. Power was maximally decreased with a phase advance of the LED command over the LFP around 90°, whilst it was maximally increased with a phase delay around 90° (Fig. 3b). The relative phases at which frequency and power were altered are however consistent with the behavior of a normal form description of a super-critical Hopf bifurcation in the vicinity of its limit-cycle. In this scenario, the LFP would approximate an observed variable, and the optogenetic drive would act in the direction of a hidden variable at a +90° angle to the LFP.

### Gamma clamp affects the timing, not rate, of pyramidal neuron firing

Although the average illumination intensity was not altered during the gamma clamp, for certain LED-LFP phase relationships gamma power increased or decreased robustly. Inhibitory currents in principal neurons, rather than spikes or excitatory currents, have previously been shown to be the main determinant of the LFP^46^, suggesting that the change in power during the clamp is not a direct effect of the optogenetic drive but results instead from a change in pyramidal neuron synchrony or phase, in a reciprocal relationship with the degree and temporal synchrony of interneuron recruitment. To determine how the clamp affects pyramidal neuron firing, we repeated experiments with an additional patch pipette to record from individual pyramidal neurons in cell-attached mode. Individual action potentials were used to align the simultaneously recorded LFP, and to estimate the phase at which they occurred. During an unmodulated ramp, pyramidal cells tended to spike sparsely, close to the trough of the oscillation, consistent with previous studies of pharmacologically induced oscillations^47^. During the clamp, an increase in oscillatory power was associated with a corresponding increase in the degree of synchrony of pyramidal cell firing: the circular dispersion of LFP phase at which pyramidal cells fired decreased relative to the unclamped situation (Fig. 4a). Conversely, a decrease in power was accompanied by a relative desynchronization of pyramidal cell firing. This relationship was qualitatively consistent, as indicated by the change in vector length obtained from the circular average of spike phases (Fig. 4c). Strikingly, however, there was no change in the overall firing rate of pyramidal cells when the oscillation power was increased or decreased by the clamp. Changes in power were thus achieved by tightening the synchrony of firing, or by desynchronizing action potentials, rather than by altering the overall activity of pyramidal neurons.

**Fig. 4.**
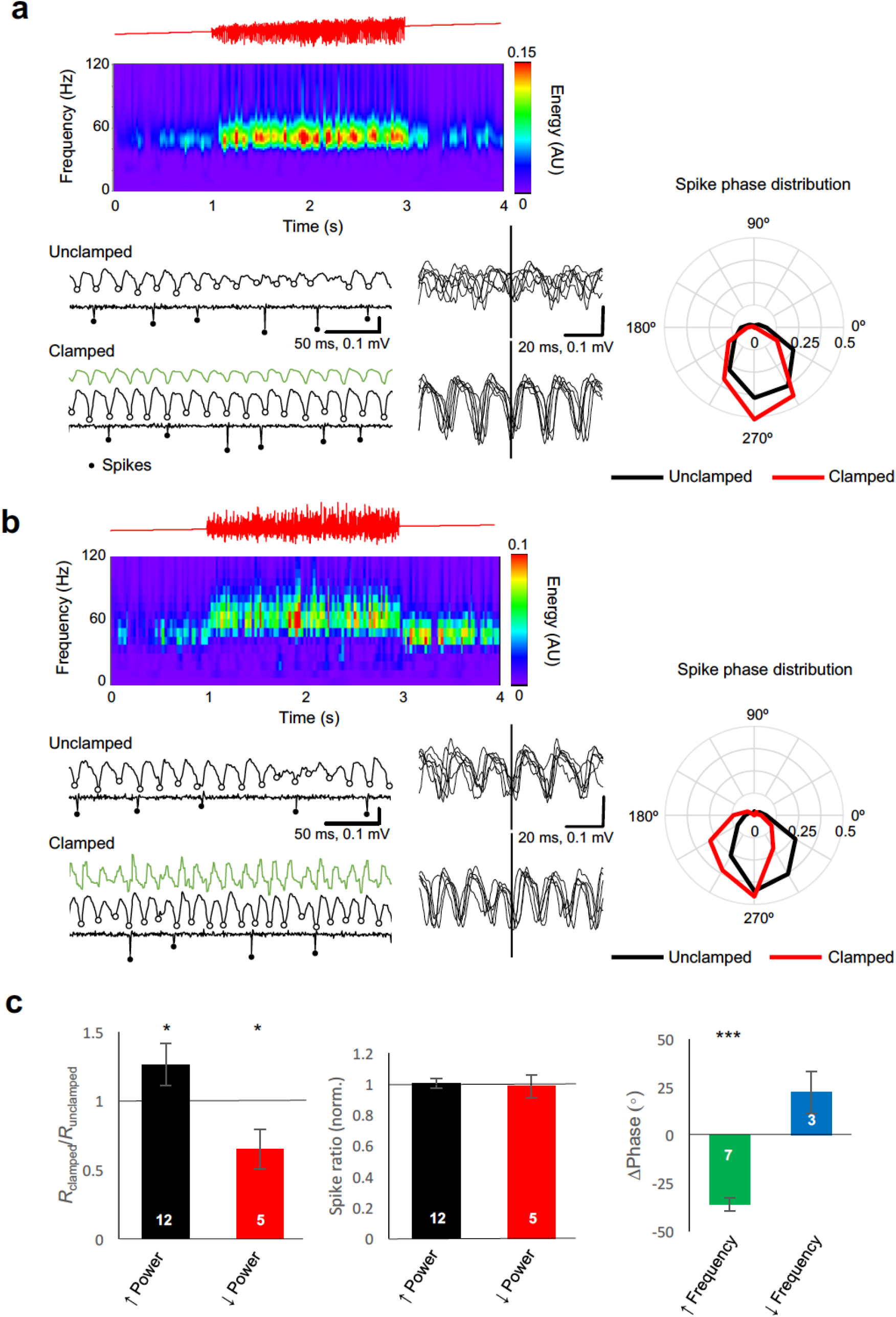
Gamma clamp alters the synchrony and phase, rather than rate, of principal cell firing. **(a)** Example closed loop modulation increasing gamma power. Top: red trace showing ramp command. Middle: spectrogram. Bottom: sample traces before (Unclamped) and during (Clamped) closed-loop modulation, showing the LFP and the cell-attached recording with identified spikes highlighted. LFP troughs are indicated by open circles. Six representative LFP traces, aligned by spike time, are shown at right. The polar plot indicates the distribution of spike phase for unclamped (black) and clamped (red) periods (averaged from 32 trials). The circular histograms sample spikes in 30° bins, and show a decrease in dispersion of spike phase during gamma clamp. **(b)** Example closed loop modulation increasing gamma frequency, plotted as for (A). The polar plot indicates phase advance of spiking. **(c)** Left: Bidirectional changes in power were associated with corresponding changes in the vector length (R) obtained by averaging all spike phases. This is consistent with a decrease in phase dispersion observed with an increase in power, and conversely, an increase in phase scatter with a decrease in power. Changes in power however did not affect the average rate of spiking, when compared with trials when gamma clamp was not applied (middle). Right: increased gamma frequency was associated with a significant phase advance of spiking. *: p<0.05; ***: p<0.001, Hotellier test^48^. Numbers of experiments are indicated in the bars.

Increases in oscillatory frequency were accompanied by a phase advance of pyramidal cell firing relative to the LFP (Fig. 4b, c). A trend for a phase delay was observed in a small number of experiments where frequency-lowering clamp was tested. This observation is consistent with the view that changes in the phase of pyramidal neuron action potentials are causally upstream of changes in gamma frequency, even though the current generators of the LFP itself are dominated by GABAergic signaling^46,49,50^.

### Excitatory current phase in principal cells determines changes in gamma spectral properties

In the examples illustrated in Figs. 4a and b, the optogenetic modulation was applied with a phase advance over the LFP of ~0° and ~45° respectively. Why does in-phase modulation result in an increase in power, and phase-advanced modulation result in an increase in frequency? To gain a mechanistic insight into how “gamma clamp” operates, we examined the phase of excitation experienced by pyramidal neurons during different clamp regimes.

We repeated experiments as above, but with one pipette used to voltage–clamp a pyramidal neuron at the estimated GABA_A_ reversal potential (approximately −70 mV), and the other pipette to record the LFP. We then measured the inward current at each phase of the gamma oscillation, as defined by the LFP, and repeated this over consecutive cycles to obtain an average time-course (Fig. 5a). The minimum (that is, least negative) inward current during the average cycle was subtracted to yield an estimate of the phasic excitatory current, which could then be represented as a vector representing its average phase and amplitude (Fig. 5b). During unclamped gamma, the excitatory current was small, and its average phase relative to the LFP varied among experiments, as expected from the very sparse synaptic connectivity among pyramidal neurons in CA1^51^. Gamma clamp imposed a large phasic inward current (Fig. 5b). Subtracting the vector representing the baseline phasic inward current yielded a vector representing the net excitatory current imposed by the gamma clamp (ΔE). This lagged behind the LED modulation, reflecting in part the opsin activation and deactivation kinetics (Fig. 5c, and arrows in Fig. 5b, right). For the example illustrated in Fig. 5 an 83° phase advance of the LED over the LFP resulted in ΔE with 113° phase delay relative to the LFP, or a total phase delay of 196° relative to the LED. This yielded an increase in frequency and decrease in power of the gamma oscillation.

**Fig. 5.**
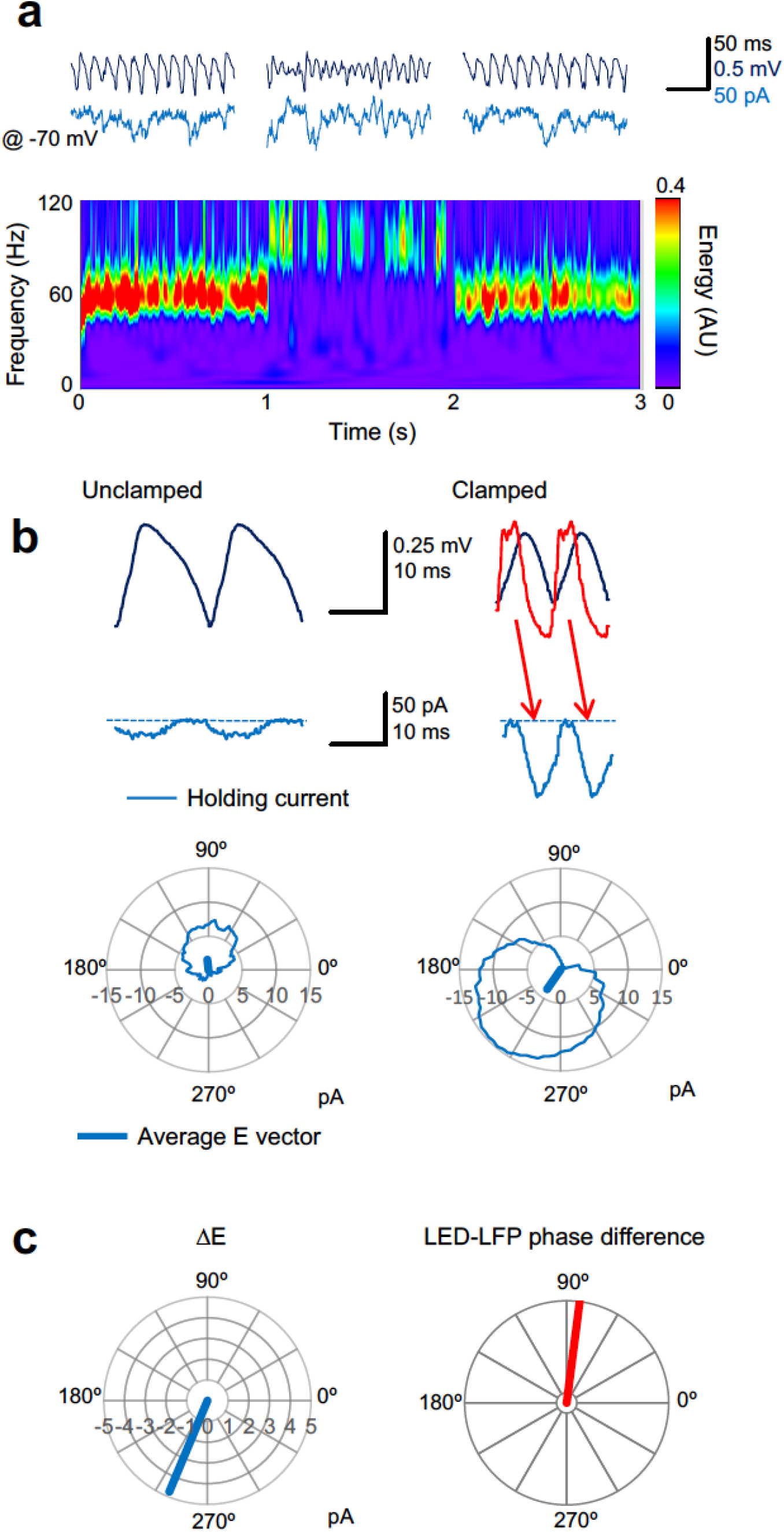
Gamma clamp imposes a phasic excitatory current to pyramidal neurons. **(a)** Top: sample LFP (black) and simultaneously recorded holding current in one pyramidal neuron held at −70 mV (blue) before, during and after feedback modulation increasing oscillatory frequency. Bottom: spectrogram. **(b)** Two cycles of the average LFP waveform and membrane current without (Unclamped) and with gamma clamp (Clamped). The average phase-advanced LED command during feedback modulation is shown superimposed (red). The minimum (least negative) inward current was subtracted (dashed lines) to estimate the phasic excitation. The red arrows indicate the temporal relationship between the peak LED driver command and the maximal excitatory current. Bottom: polar plots indicating the cycle-average of the excitatory current during unclamped (left) and clamped (right) periods of the trial shown in (a). The vectors indicate the average phases of the currents. **(c)** Left: difference vector obtained from the vectors in (b), representing the net phasic excitatory current imposed by gamma clamp. Right: phase difference between LED and LFP for the same experiment.

Comparing across different clamp regimes reveals how gamma frequency and power change in relation to the phasic excitation experienced by principal cells (Fig. 6a, b). An increase in gamma frequency was achieved when the average excitatory current phase occurred during the down-stroke and trough of the LFP (~200 to 300°), whilst a decrease in frequency was achieved when excitation was applied around the peak of the LFP (~70°). A maximal increase in power, on the other hand, was achieved with excitation around 20 °, coinciding with the upstroke of the LFP, and a decrease in power occurred with excitation around 200°. Taking into account that, under baseline conditions, pyramidal neurons fire maximally around 285°, these data imply that the increase in frequency occurs because they are brought to firing threshold earlier (see also Fig. 4b, c). An increase in power, on the other hand, occurs because pyramidal neurons are synchronized by adding a depolarization shortly after they have fired (see also Fig. 4a, c). Conversely, the oscillation frequency decreases when phasic depolarization is reduced as pyramidal neurons approach their maximal firing probability.

**Fig. 6.**
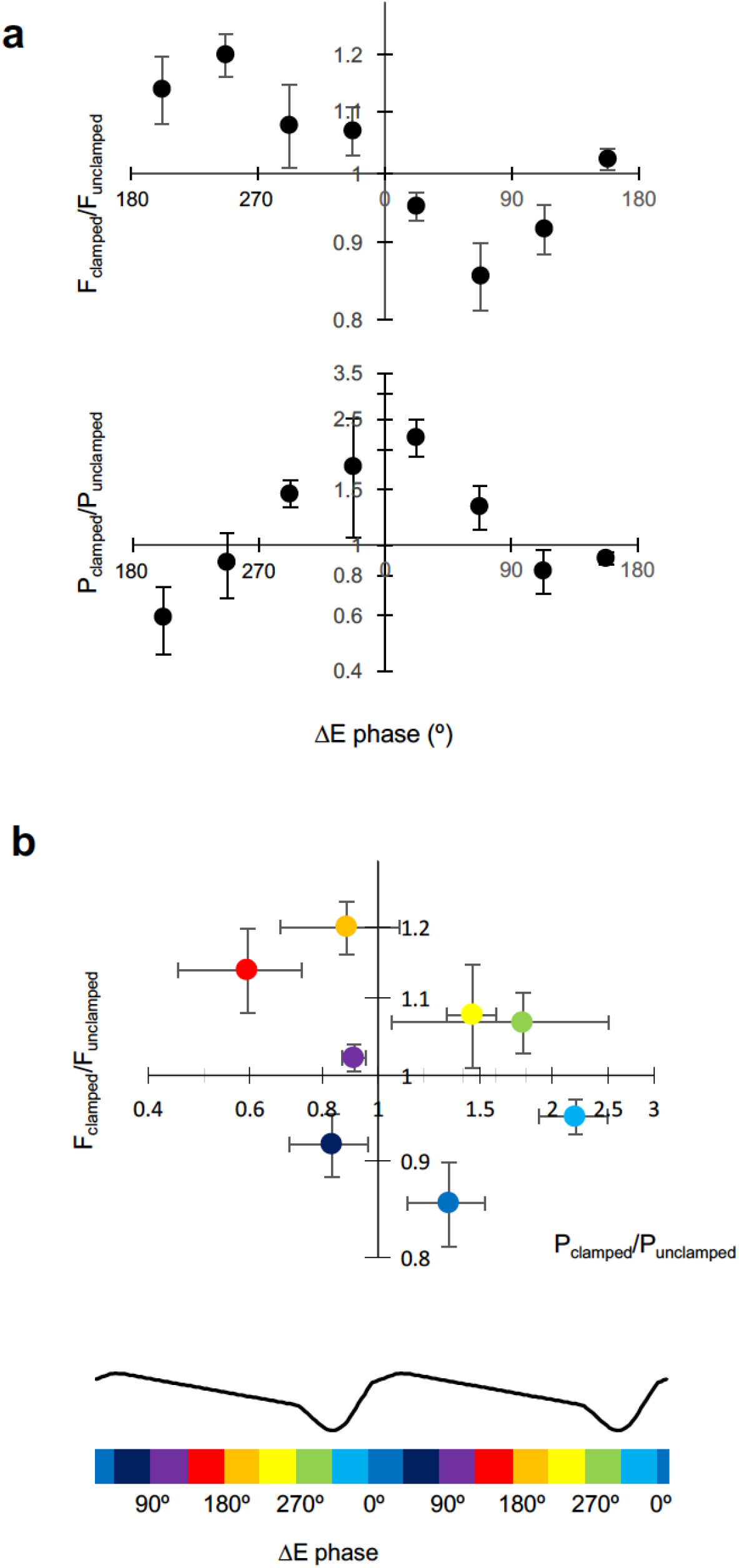
Excitatory current phase determines changes in frequency and power. **(a)** Change in gamma frequency and power, plotted against the phase of the net excitatory current (ΔE) calculated as in Fig. 5. **(b)** F_clamped_/F_unclamped_ plotted against P_clamped_/P_unclamped_ for different excitatory current phases, indicated by the color code below, aligned with the average LFP waveform. Pyramidal neurons spike around 285°.

## Discussion

The present study shows that closed-loop optogenetic manipulation of principal cells allows predictable, bidirectional and dissociable changes in the power and frequency of gamma oscillations. We observed a broad consistency between the frequency manipulation achieved with closed loop optogenetic feedback and that predicted from the phase response behavior previously observed with intermittent optogenetic stimuli^10^. Optogenetically and pharmacologically induced gamma also exhibited similar dynamical properties in that study, implying that the principles uncovered in the present work are not specific to the way gamma oscillations were elicited.

Previous studies have stressed the importance of fast-spiking parvalbumin-positive (PV+) interneurons in gamma^20,21^ (but see ref. ^36^). PV+ basket cells tend to fire with very little phase dispersion, close to one-to-one with each cycle of the oscillation *in vitro*^17,49^. Our attempts to achieve gamma clamp by targeting interneurons rather than pyramidal cells have thus far been unsuccessful because their out of phase recruitment powerfully suppresses the oscillation (data not shown). The weaker phase-locking of pyramidal than PV+ cell firing to gamma oscillations, together with their sparse firing on successive cycles of gamma^49,52,53^, may however confer a broader dynamic range over which they can influence the phase, frequency and amplitude of the oscillation. Taken together with previous evidence that open-loop sinusoidal optogenetic stimulation of principal cells can entrain a gamma oscillation^10^, the present data underline the importance of action potential timing in principal cells in the spectral and temporal properties of hippocampal gamma, notwithstanding the evidence that the LFP itself is dominated by inhibitory currents in principal cells^46^, and argue against a model where the function of principal cells is only to depolarize a population of reciprocally connected interneurons.

Closed-loop manipulations have been applied previously in the context of network oscillations, using either electrical and optogenetic stimuli delivered at specific phases of theta or gamma oscillations, in order to probe the mechanisms of long-term plasticity induction^54,55^ or sharp-wave ripple generation^33^, or to test the theta phase-dependence of memory encoding and retrieval^34^. A similar strategy has been used to interrupt experimental thalamocortical seizures^56^. However, these studies have not aimed at modulating the amplitude or frequency of an on-going oscillation.

We have focused on gamma because a local circuit is sufficient to generate the oscillation, and we have previously shown that the phase response behavior of hippocampal gamma is well described by a simple dynamical model^10^. The circuits underlying theta and other oscillations either involve longer-range connections in the brain or are poorly defined. They are therefore less likely to be amenable to local optogenetic manipulation. This does not exclude the possibility that, for instance, theta oscillations in the hippocampus could be manipulated by closed-loop modulation of excitability in the basal forebrain.

The ability to alter the amplitude and frequency of gamma suggests a versatile tool to test the roles of gamma in information routing and other high-level brain functions, both in health and in disease states such as schizophrenia^57^. Hitherto, most experimental manipulations of oscillations have relied on periodic stimulation, which can entrain network oscillations^10^ or evoke oscillations in an otherwise asynchronous network^20,21^. Transcranial stimulation designed to entrain oscillations in vivo can bias perception^58–60^ and bidirectionally affect performance in motor^61^ and working memory^62^ tasks. However, external periodic stimulation is not well suited to desynchronize network activity or to suppress oscillatory dynamics. Furthermore, if periodic stimulation is used, the desired change in amplitude or frequency is achieved at the cost of imposing an externally determined phase on the oscillation. This will prevent the oscillation from entraining to endogenous periodic signals such as those arising from other oscillating networks or periodic sensory stimuli.

Closed-loop stimulation, in which signals recorded from a network are used in real time to bias its state, in principle provides an alternative way of manipulating network oscillations, and has been used to interfere with pathological rhythms in models of Parkinson’s disease^63^, to suppress Parkinsonian tremor^64^, and in a model of thalamocortical epilepsy^65^. This approach relies on an artificial feedback loop which either counteracts or amplifies the endogenous feedback responsible for synchronizing the network^66^. Importantly, optogenetics has the advantage over electrical stimulation that the modulation can be distributed across a population of neurons. We have, moreover, shown that closed-loop manipulation of a gamma oscillation can be achieved without a net increase or decrease in the average firing rate of neurons, implying that it would not necessarily perturb information represented as an average firing rate code.

Extrapolating from *in vitro* gamma to the brain *in situ* presents several technical challenges, including the need for optical fibers to illuminate the tissue and the potential for photoelectrical artifacts. Moreover, oscillations are generally less prominent because the current generators from multiple oscillating and non-oscillating populations overlap, complicating the evaluation of phase and frequency. Nevertheless, the present study identifies some general principles to guide attempts to achieve bidirectional and dissociable modulation of oscillatory frequency and power *in vivo*. This should allow a definitive test of the causal role of gamma in functions such as attention modulation and information routing^67^.

## Online Methods

All procedures followed the Animals (Scientific Procedures) Act, 1986. P21 male C57 mice were anesthetized with isoflurane and placed in a stereotaxic frame (Kopf Instruments). A suspension of AAV5-CaMKIIα-C1V1(E122T/E162T)-TS-eYFP (UNC Vector Core, titer 5 × 10^12^ IU/ml) was injected at a rate of 100 nl/min into 4 sites in both hippocampi (injection volume: 300-500 nl per site). The antero-posterior injection coordinate was taken as 2/3 of the distance from bregma to lambda. The lateral coordinates were 3.0 mm from the midline, and the ventral coordinates were 3.5, 3.0, 2.5 and 2.0 mm from the surface of the skull.

Hippocampal slices were prepared at least 4 weeks later. Animals were sacrificed by pentobarbitone overdose and underwent transcardiac perfusion with an oxygenated solution containing (in mM): 92 N-methyl-D-glucamine-Cl, 2.5 KCl, 1.25 NaH_2_PO_4_, 20 HEPES, 30 NaHCO_3_, 25 glucose, 10 MgCl_2_, 0.5 CaCl_2_, 2 thiourea, 5 Na-ascorbate and 3 Na-pyruvate, with sucrose added to achieve an osmolality of 315 mOsm/L. Brain slices (400 μm thick) were prepared at room temperature and then incubated at 37 °C for 12 minutes in the same solution. They were subsequently stored at room temperature, in a solution containing (in mM): 126 NaCl, 3 KCl, 1.25 NaH_2_PO_4_, 2 MgSO_4_, 2 CaCl_2_, 24 NaHCO_3_, 10 glucose, shielded from light, before being transferred to the stage of an upright microscope (Olympus BX51WI or Scientifica SliceScope), where they were perfused on both sides with the same solution at 32° C. Expression of C1V1 in CA1 was verified by epifluorescence, and CA3 was ablated to focus on local gamma-generating mechanisms.

Epifluorescence imaging and C1V1 stimulation were achieved with LEDs (OptoLED, Cairn Instruments, or assembled from Thorlabs components using an M590L2 590 nm LED and a DC2100 high-power LED driver). The light source was coupled to the epifluorescence illuminator of the microscope, with a silver mirror in the place of a dichroic cube. Wide-field illumination was delivered via a 20x, 0.5 NA water immersion objective. The current delivered to the LED was kept in the linear input-output range, and the irradiance was <5 mW/mm^2^. Light ramps typically lasting 8 s were delivered every 30 – 45 s.

LFPs were recorded in the CA1 pyramidal layer using patch pipettes filled with extracellular solution and a Multiclamp 700B amplifier (Molecular Devices), and band-pass filtered between 1 and 200 or 500 Hz. A linear LED ramp command was generated via a multifunction data acquisition card (National Instruments PCI-6221) and, together with the LFP, was digitized using a real-time controller (National Instruments cRIO-9022) with a Xilinx Virtex-5 FPGA (cRIO-9133) operating at a loop rate of 10 kHz. The ramp was multiplied by (1 + *k*_1_LFP + *k*_2_dLFP/*d*t), stepping through different values of *k* in a pseudo-random order for successive trials. dLFP/dt was calculated as the difference between successive digitization values in the FPGA, averaged over successive 2 ms intervals to minimize high-frequency noise. The output of the FPGA/real-time controller was sent to the LED driver, and digitized in parallel with the LFP at 10 kHz on the data acquisition PC.

To study the phase relationship of action potentials and the LFP oscillation, a cell-attached recording was obtained using a second patch pipette held in voltage clamp mode, low-pass filtered at 10 kHz and digitized in parallel with the LFP and LED command signal. The phasic excitatory current was recorded in the same way, but using a whole-cell pipette containing (in mM): K-gluconate (145), NaCl (8), KOH-HEPES (10), EGTA (0.2), Mg-ATP (2) and Na 3 - GTP (0.3); pH 7.2; 290 mOsm.

Off-line analysis was performed in LabVIEW (National Instruments) and R. Time-frequency spectrograms were calculated using a Morlet wavelet transform and are displayed as heat maps. Because the gamma oscillation was non-stationary, its frequency was estimated by calculating the short-term Fourier transform and then averaging the mean instantaneous frequency for successive overlapping intervals. The power of the oscillation was estimated in the same way, by averaging the power at the mean instantaneous frequency.

Spikes were identified using threshold crossing. The instantaneous oscillation phase was estimated by passing a 200-ms segment of the LFP centered on the spike through a Hanning window, and then calculating its phase and frequency using the Extract Single Tone VI in LabVIEW.

To estimate the phase relationship between spikes or excitatory currents and the gamma oscillation, we first identified successive troughs of the LFP using the WA Multiscale Peak Detection VI in LabVIEW. Gamma cycles that deviated more than 20% from the modal period were rejected. The membrane current waveform between successive troughs was then expressed as a function of instantaneous phase and averaged over all accepted cycles in the interval. The minimal (least negative) inward current was subtracted to yield the average phasic current waveform.

## Acknowledgments

We are grateful to Francis Carpenter, Flora Lee and Iris Oren for pilot experiments. This work was supported by the Wellcome Trust.

## References

1. Akam, T. & Kullmann, D. M. Oscillations and filtering networks support flexible routing of information. Neuron 67, 308–320 (2010).

2. Akam, T. & Kullmann, D. M. Oscillatory multiplexing of population codes for selective communication in the mammalian brain. Nat. Rev. Neurosci. 15, 111–122 (2014).

3. Fries, P. A mechanism for cognitive dynamics: neuronal communication through neuronal coherence. Trends Cogn. Sci. 9, 474–80 (2005).

4. Lisman, J. Working memory: the importance of theta and gamma oscillations. Curr. Biol. CB 20, R490–492 (2010).

5. Rodriguez, E. et al. Perception’s shadow: long-distance synchronization of human brain activity. Nature 397, 430–433 (1999).

6. Salinas, E. & Sejnowski, T. J. Correlated neuronal activity and the flow of neural information. Nat. Rev. Neurosci. 2, 539–550 (2001).

7. Schnitzler, A. & Gross, J. Normal and pathological oscillatory communication in the brain. Nat Rev Neurosci 6, 285–296 (2005).

8. Börgers, C. & Kopell, N. Synchronization in networks of excitatory and inhibitory neurons with sparse, random connectivity. Neural Comput. 15, 509–538 (2003).

9. Kirst, C., Timme, M. & Battaglia, D. Dynamic information routing in complex networks. Nat. Commun. 7, 11061 (2016).

10. Akam, T., Oren, I., Mantoan, L., Ferenczi, E. & Kullmann, D. M. Oscillatory dynamics in the hippocampus support dentate gyrus–CA3 coupling. Nat. Neurosci. 15, 763–768 (2012).

11. Fries, P. Rhythms for Cognition: Communication through Coherence. Neuron 88, 220–235 (2015).

12. Varela, F., Lachaux, J. P., Rodriguez, E. & Martinerie, J. The brainweb: phase synchronization and large-scale integration. Nat. Rev. Neurosci. 2, 229–239 (2001).

13. Bosman, C. A. et al. Attentional stimulus selection through selective synchronization between monkey visual areas. Neuron 75, 875–888 (2012).

14. Grothe, I., Neitzel, S. D., Mandon, S. & Kreiter, A. K. Switching neuronal inputs by differential modulations of gamma-band phase-coherence. J. Neurosci. 32, 16172–16180 (2012).

15. Colgin, L. L. et al. Frequency of gamma oscillations routes flow of information in the hippocampus. Nature 462, 353–357 (2009).

16. Buzsáki, G. & Schomburg, E. W. What does gamma coherence tell us about inter-regional neural communication? Nat. Neurosci. 18, 484–489 (2015).

17. Bartos, M., Vida, I. & Jonas, P. Synaptic mechanisms of synchronized gamma oscillations in inhibitory interneuron networks. Nat. Rev. Neurosci. 8, 45–56 (2007).

18. Buzsáki, G. & Wang, X.-J. Mechanisms of gamma oscillations. Annu. Rev. Neurosci. 35, 203–225 (2012).

19. Whittington, M. A., Traub, R. D. & Jefferys, J. G. Synchronized oscillations in interneuron networks driven by metabotropic glutamate receptor activation. Nature 373, 612–615 (1995).

20. Cardin, J. A. et al. Driving fast-spiking cells induces gamma rhythm and controls sensory responses. Nature 459, 663–667 (2009).

21. Sohal, V. S., Zhang, F., Yizhar, O. & Deisseroth, K. Parvalbumin neurons and gamma rhythms enhance cortical circuit performance. Nature 459, 698–702 (2009).

22. Mann, E. O., Radcliffe, C. A. & Paulsen, O. Hippocampal gamma-frequency oscillations: from interneurones to pyramidal cells, and back. J. Physiol. 562, 55–63 (2005).

23. Bartos, M. et al. Fast synaptic inhibition promotes synchronized gamma oscillations in hippocampal interneuron networks. Proc. Natl. Acad. Sci. U. S. A. 99, 13222–7 (2002).

24. Fisahn, A. et al. Distinct roles for the kainate receptor subunits GluR5 and GluR6 in kainate-induced hippocampal gamma oscillations. J. Neurosci. 24, 9658–9668 (2004).

25. Wang, X. J. & Buzsáki, G. Gamma oscillation by synaptic inhibition in a hippocampal interneuronal network model. J. Neurosci. 16, 6402–6413 (1996).

26. Tiesinga, P. & Sejnowski, T. J. Cortical Enlightenment: Are Attentional Gamma Oscillations Driven by ING or PING? Neuron 63, 727–732 (2009).

27. Wilson, H. R. & Cowan, J. D. Excitatory and inhibitory interactions in localized populations of model neurons. Biophys. J. 12, 1–24 (1972).

28. Akao, A., Ogawa, Y., Jimbo, Y., Ermentrout, G. B. & Kotani, K. Relationship between the mechanisms of gamma rhythm generation and the magnitude of the macroscopic phase response function in a population of excitatory and inhibitory modified quadratic integrate-and-fire neurons. Phys. Rev. E 97, 012209 (2018).

29. Kotani, K., Yamaguchi, I., Yoshida, L., Jimbo, Y. & Ermentrout, G. B. Population dynamics of the modified theta model: macroscopic phase reduction and bifurcation analysis link microscopic neuronal interactions to macroscopic gamma oscillation. J. R. Soc. Interface 11, 20140058 (2014).

30. Grosenick, L., Marshel, J. H. & Deisseroth, K. Closed-loop and activity-guided optogenetic control. Neuron 86, 106–139 (2015).

31. Krook-Magnuson, E., Armstrong, C., Oijala, M. & Soltesz, I. On-demand optogenetic control of spontaneous seizures in temporal lobe epilepsy. Nat. Commun. 4, 1376 (2013).

32. Paz, J. T. et al. Closed-loop optogenetic control of thalamus as a tool for interrupting seizures after cortical injury. Nat. Neurosci. 16, 64–70 (2013).

33. Stark, E. et al. Pyramidal cell-interneuron interactions underlie hippocampal ripple oscillations. Neuron 83, 467–480 (2014).

34. Siegle, J. H. & Wilson, M. A. Enhancement of encoding and retrieval functions through theta phase-specific manipulation of hippocampus. eLife 3, e03061 (2014).

35. Newman, J. P. et al. Optogenetic feedback control of neural activity. eLife 4, e07192 (2015).

36. Veit, J., Hakim, R., Jadi, M. P., Sejnowski, T. J. & Adesnik, H. Cortical gamma band synchronization through somatostatin interneurons. Nat. Neurosci. 20, 951–959 (2017).

37. Witt, A. et al. Controlling the oscillation phase through precisely timed closed-loop optogenetic stimulation: a computational study. Front. Neural Circuits 7, (2013).

38. Yizhar, O. et al. Neocortical excitation/inhibition balance in information processing and social dysfunction. Nature 477, 171–178 (2011).

39. Adesnik, H. Layer-specific excitation/inhibition balances during neuronal synchronization in the visual cortex. J. Physiol. (2018). doi:10.1113/JP274986

40. Adesnik, H. & Scanziani, M. Lateral competition for cortical space by layer-specific horizontal circuits. Nature 464, 1155–1160 (2010).

41. Butler, J. L., Mendonça, P. R. F., Robinson, H. P. C. & Paulsen, O. Intrinsic Cornu Ammonis Area 1 Theta-Nested Gamma Oscillations Induced by Optogenetic Theta Frequency Stimulation. J. Neurosci. 36, 4155–4169 (2016).

42. Pastoll, H., Solanka, L., van Rossum, M. C. W. & Nolan, M. F. Feedback inhibition enables θ-nested γ oscillations and grid firing fields. Neuron 77, 141–154 (2013).

43. Ni, J. et al. Gamma-Rhythmic Gain Modulation. Neuron 92, 240–251 (2016).

44. Lu, Y. et al. Optogenetically induced spatiotemporal gamma oscillations and neuronal spiking activity in primate motor cortex. J. Neurophysiol. 113, 3574–3587 (2015).

45. Cole, S. R. & Voytek, B. Brain Oscillations and the Importance of Waveform Shape. Trends Cogn. Sci. 21, 137–149 (2017).

46. Oren, I., Hájos, N. & Paulsen, O. Identification of the current generator underlying cholinergically induced gamma frequency field potential oscillations in the hippocampal CA3 region. J. Physiol. 588, 785–797 (2010).

47. Fisahn, A., Pike, F. G., Buhl, E. H. & Paulsen, O. Cholinergic induction of network oscillations at 40Hz in the hippocampus in vitro. Nature 394, 186–189 (1998).

48. Zar, J. H. Biostatistical Analysis 5th By Jerrold H. Zar. (PIE, 2009).

49. Gulyás, A. I. et al. Parvalbumin-containing fast-spiking basket cells generate the field potential oscillations induced by cholinergic receptor activation in the hippocampus. J. Neurosci. 30, 15134–15145 (2010).

50. Hájos, N. et al. Spike Timing of Distinct Types of GABAergic Interneuron during Hippocampal Gamma Oscillations In Vitro. J. Neurosci. 24, 9127–9137 (2004).

51. Deuchars, J. & Thomson, A. M. CA1 pyramid-pyramid connections in rat hippocampus in vitro: dual intracellular recordings with biocytin filling. Neuroscience 74, 1009–1018 (1996).

52. Tukker, J. J., Fuentealba, P., Hartwich, K., Somogyi, P. & Klausberger, T. Cell type-specific tuning of hippocampal interneuron firing during gamma oscillations in vivo. J. Neurosci. 27, 8184–9 (2007).

53. Csicsvari, J., Jamieson, B., Wise, K. D. & Buzsáki, G. Mechanisms of gamma oscillations in the hippocampus of the behaving rat. Neuron 37, 311–322 (2003).

54. Huerta, P. T. & Lisman, J. E. Bidirectional synaptic plasticity induced by a single burst during cholinergic theta oscillation in CA1 in vitro. Neuron 15, 1053–1063 (1995).

55. Pavlides, C., Greenstein, Y. J., Grudman, M. & Winson, J. Long-term potentiation in the dentate gyrus is induced preferentially on the positive phase of theta-rhythm. Brain Res. 439, 383–387 (1988).

56. Berényi, A., Belluscio, M., Mao, D. & Buzsáki, G. Closed-Loop Control of Epilepsy by Transcranial Electrical Stimulation. Science 337, 735–737 (2012).

57. Uhlhaas, P. J. & Singer, W. Abnormal neural oscillations and synchrony in schizophrenia. Nat. Rev. Neurosci. 11, 100–113 (2010).

58. Neuling, T., Rach, S., Wagner, S., Wolters, C. H. & Herrmann, C. S. Good vibrations: oscillatory phase shapes perception. NeuroImage 63, 771–778 (2012).

59. Thut, G., Schyns, P. G. & Gross, J. Entrainment of perceptually relevant brain oscillations by non-invasive rhythmic stimulation of the human brain. Front. Psychol. 2, 170 (2011).

60. Romei, V., Gross, J. & Thut, G. On the role of prestimulus alpha rhythms over occipitoparietal areas in visual input regulation: correlation or causation? J. Neurosci. 30, 8692–8697 (2010).

61. Joundi, R. A., Jenkinson, N., Brittain, J.-S., Aziz, T. Z. & Brown, P. Driving Oscillatory Activity in the Human Cortex Enhances Motor Performance. Curr. Biol. 22, 403–407 (2012).

62. Polanía, R., Nitsche, M. A., Korman, C., Batsikadze, G. & Paulus, W. The importance of timing in segregated theta phase-coupling for cognitive performance. Curr. Biol. CB 22, 1314–1318 (2012).

63. Rosin, B. et al. Closed-Loop Deep Brain Stimulation Is Superior in Ameliorating Parkinsonism. Neuron 72, 370–384 (2011).

64. Brittain, J.-S., Probert-Smith, P., Aziz, T. Z. & Brown, P. Tremor suppression by rhythmic transcranial current stimulation. Curr. Biol. CB 23, 436–440 (2013).

65. Butt, S. J. B. et al. The temporal and spatial origins of cortical interneurons predict their physiological subtype. Neuron 48, 591–604 (2005).

66. Rosenblum, M. G. & Pikovsky, A. S. Controlling synchronization in an ensemble of globally coupled oscillators. Phys. Rev. Lett. 92, 114102 (2004).

67. Sohal, V. S. How Close Are We to Understanding What (if Anything) γ Oscillations Do in Cortical Circuits? J. Neurosci. 36, 10489–10495 (2016).

